# Early-life microbiota disruption by antibiotics elicits fitness trade-offs that differ by sex

**DOI:** 10.1101/2025.08.26.670495

**Authors:** Laura D. Schell, Grace Rubin, Eric Chan, Rachel N. Carmody

## Abstract

Exposure to early-life antibiotics (ELA) promotes adult obesity across diverse species, often more strongly in males than females^1–3^. However, the physiological and evolutionary mechanisms driving ELA-mediated developmental plasticity and their consequences for fitness remain unclear^3^. Here, treating young mice with high– or low-dose ampicillin altered the gut microbiome and elicited reduced lean mass and energy expenditure in males to promote increased adult visceral adiposity. High-dose ELA constrained energy availability during treatment, evidenced by reduced growth and cecal short-chain fatty acids, raising the possibility that developmental plasticity under ELA could be adaptive in energy-limited environments. Adult ELA-treated males had reduced fitness under free-fed conditions — driven by smaller body size, fat stores, and impaired immunity — but were buffered against fitness reductions under caloric restriction, consistent with predictive adaptive response models of development^4,5^. Strikingly, ELA-treated females exhibited none of the proximate metabolic responses observed in males, with adult body size and composition indistinguishable from controls. In the absence of similar developmental plasticity, ELA-treated females exhibited exacerbated fitness reductions under caloric restriction, including lower energetic investments in reproduction and immunity. Our results suggest that ELA exposure elicits sexually dimorphic developmental responses that engender long-term health and fitness consequences dependent on adult nutritional conditions.

## INTRODUCTION

Exposure to antibiotics early in life can promote metabolic and immune-related diseases in adulthood. In humans, antibiotic use in infancy is associated with increased incidence of adult obesity, allergy/asthma, and inflammatory bowel disease^1,2,6–8^. Similarly, young mice treated with pulsed therapeutic or chronic subtherapeutic antibiotics exhibit greater levels of body fat, glucose intolerance, allergy/asthma, intestinal inflammation, and increased susceptibility to infections in adulthood^8–13^. These phenotypes suggest that early-life antibiotic (ELA) treatment perturbs metabolic and immune development. Here, we explore how and why ELA treatment alters development, evaluating alternative strategies of energy allocation and their consequences for fitness. In doing so, we test the extent to which ELA-mediated developmental plasticity could be considered an adaptive response to early-life adversity.

Studies of ELA-induced obesity have demonstrated that both pulsed high-dose (therapeutic) and chronic low-dose (subtherapeutic) exposures disrupt the infant gut microbiome and result in adult adiposity, even without increased caloric intake^3,14^. These outcomes suggest that ELA exposure increases dietary energy harvest and/or prompts energetic trade-offs that increase allocation of available calories to fat storage versus other physiological investments like growth, immune function, reproduction, or physical activity. The gut microbiome can regulate the energy balance of its host through diverse mechanisms^15^. Murine studies of low-dose ELA-induced obesity highlight several possible gut microbiome-mediated proximate mechanisms, including increased short-chain fatty acid (SCFA) production^16^, altered intestinal fat metabolism^11^, reduced bile acid metabolism by gut microbes^17^, and loss of keystone developmental taxa^9^. In the case of high-dose ELA, experiments in germ-free, fiber-deprived, and GPR41^-/-^ or GPR43^-/-^ mutant mice implicate the loss of SCFA signaling as a developmental trigger for adult obesity^3,18^.

The ultimate reasons why ELA treatment should promote host developmental plasticity at all are likewise unclear, given that antibiotics are generally indented to target microbes rather than host tissues. One possibility is that ELA-mediated developmental plasticity reflects a response to volatility in gut microbiome-mediated pathways of energy metabolism^3^. This is particularly pertinent to therapeutic, high-dose antibiotics, which decimate absolute abundance of gut microbes, impair microbiome-mediated dietary energy harvest by reducing SCFA production, and thereby promote short-term negative energy balance^18,19^. In humans, early-life energy constraint increases adult risk of metabolic disease, best evidenced by epidemiological outcomes of famines including the Dutch Hunger Winter^20^ and Ukrainian Holodomor^21^. Periods of undernutrition may be followed by periods of faster “catch-up” growth when nutritional environments improve, but adults exposed to such early-life stresses tend to attain lower adult height and have smaller internal organs^22,23^. Lower energy expenditure and metabolic capacity then predispose these individuals to obesity and type 2 diabetes (T2D) in energy-rich conditions^23^, such as in industrialized societies with ready access to highly processed, nutrient-dense foods. ELA-induced adult adiposity may therefore arise due to perturbations of host-microbiome interactions in energy metabolism that impose energy constraint^3^.

Several evolutionary models have proposed that developmental plasticity prompted by early-life energy constraint could be adaptive even if it increases the risk of metabolic disease in energy-rich adult environments. These models argue that changes in growth rate and the development of smaller, less costly bodies ultimately arise either to ensure survival during the adverse early-life environment [developmental constraints (DC) model]^24,25^, to adapt to an anticipated adverse adult environment [predictive adaptive response (PAR) model]^4,5^, or as a maternal strategy to optimize lifetime reproductive success at the cost of the current offspring’s fitness [maternal capital (MC) model]^26^. All three of these models predict that an energy-poor early-life environment combined with an energy-rich adult environment will result in increased risk of metabolic diseases. However, when early-life adversity is followed by an energy-poor adult environment, PAR predicts that individuals will be less negatively impacted than those with more optimal early-life conditions, whereas DC and MC predict exacerbated negative impacts. Interrogating these differential predictions enables inferences about the potential ultimate causes of ELA-induced developmental plasticity.

Where does the energy fueling adult adiposity after ELA come from? How do these trade-offs vary with host characteristics and antibiotic dose? Are there fitness benefits associated with ELA-induced developmental plasticity, and if so, what is the most parsimonious adaptive mechanism? Here we address these questions, interrogating the overarching hypothesis that ELA exposure is a form of early-life adversity analogous to that of variable energy availability, prompting developmental changes that serve to buffer individual fitness.

## RESULTS

### ELA promotes visceral adiposity in males, but not females, at the cost of lean body mass

We employed a previously established mouse model of ELA-induced obesity^9,10,14^, administering ampicillin via drinking water to pregnant mice and their resulting offspring. Recipients of low-dose ELA were treated with ampicillin (6.7mg/L) continuously from day 13.5 of gestation through either 4 weeks of age (L4) or until endpoint at 30 weeks (L30) (**Fig. 1A**). Recipients of high-dose ELA (HD) were treated in pulses, with ampicillin (333mg/L) administered for a week during gestation (E13.5–post-natal day 1) and for a second week immediately following weaning (weeks 3-4 of age).

**Figure 1.**
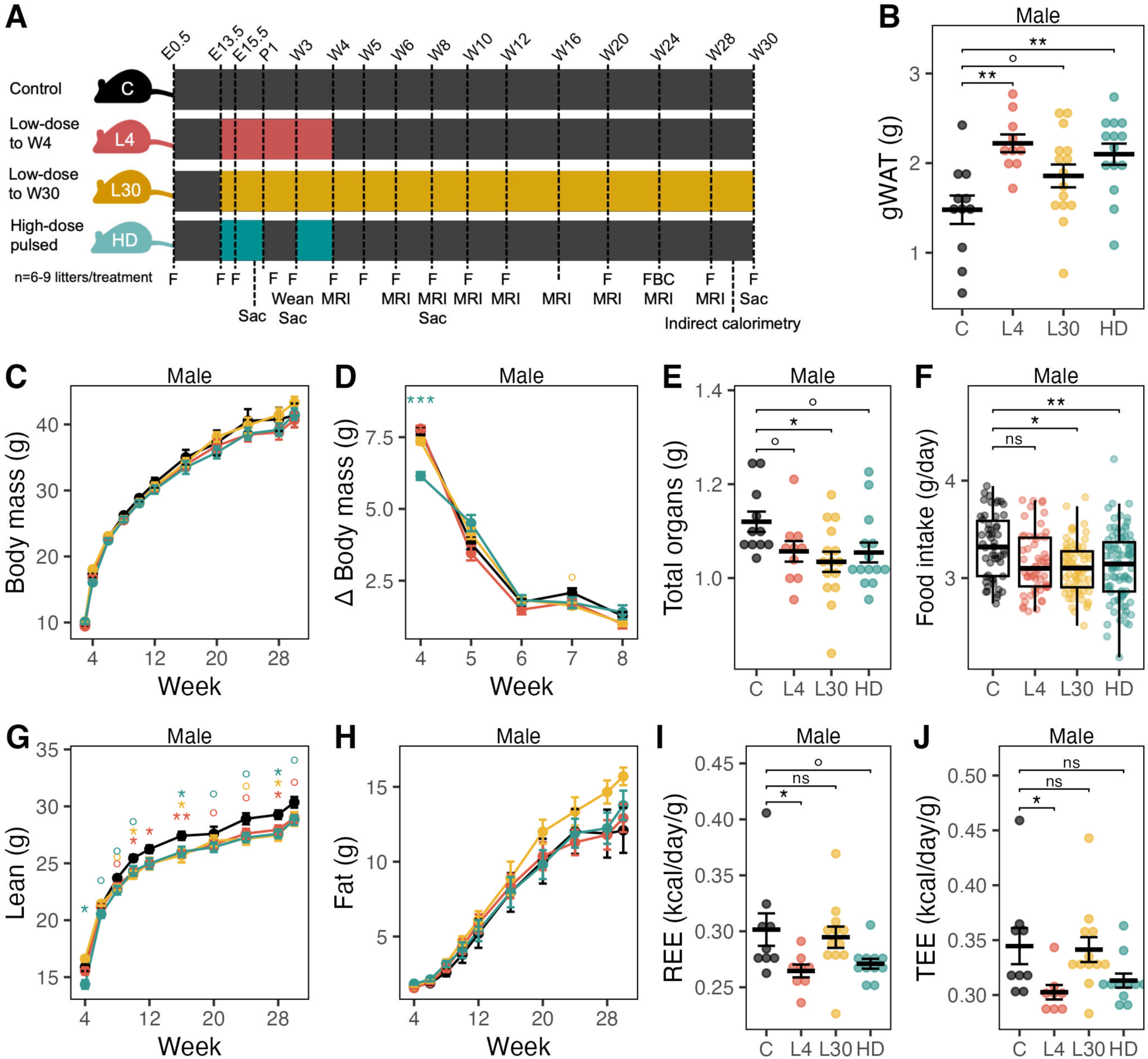
Low– and high-dose ELA treatment promotes visceral adiposity at the cost of lean mass in males. (**A**) Experimental design, from embryonic/gestational day 0.5 (E0.5) to post-natal day 1 (P1), to week 30 (W30). F = fecal sampling, MRI = body composition mesurement via EchoMRI, FBC = fecal bomb calorimetry, Sac = sacrifice and tissue collection. (**B**) Mass of gonadal (epididymal) white adipose tissue (gWAT), a visceral fat deposit, at 30 weeks of age. (**C**) Body mass from 3 to 30 weeks. (**D**). Change in body mass from 3-8 weeks, with the y-axis value representing change observed at the week shown on the x-axis relative to the previous week. (**E**) Sum of organ masses (heart, spleen, kidney, testes, brain) at 30 weeks. (**F**) Mean daily food intake per mouse. (**G**) Lean body mass and (**H**) fat mass from 4 to 30 weeks, as measured by EchoMRI. (**I**) Resting energy expenditure and (**J**) total energy expenditure, as measured by indirect calorimetry. Data are mean ± s.e.m. Wilcoxon rank-sum test with control as reference group (B-E, G-J) or linear mixed effects model (F) of Food intake ∼ Treatment + (1|Cage_ID) + (1|Timepoint), ns = p≥0.1, ° = p<0.1, * = p<0.05, ** = p<0.01, *** = p<0.001.

By 30 weeks of age, L4, L30, and HD males all had larger visceral fat deposits compared to sex-matched untreated controls (CON) (**Fig. 1B**), with similar but weaker trends in subcutaneous deposits (**Extended Data Fig. 1A**) and endpoint total body fat as indexed by EchoMRI (**Fig. 1H**). Notably, ELA-treated males did not exhibit differences in body mass compared with controls, reflecting a trade-off with lean mass, which was reduced compared with controls as early as the first measurement at 4 weeks (HD) or 8 weeks (L4 and L30) (**Fig. 1C,G**). The combined mass of internal organs was also lower across all groups of ELA males compared with controls (**Fig. 1E**). Whereas L4 and L30 mice gained body mass and lean mass at a rate similar to controls, HD mice exhibited both reduced lean mass at 3 weeks and reduced growth rate from 3-4 weeks (**Fig. 1D,G**), suggesting that high-dose ELA may constrain energy availability during treatment.

Despite higher adiposity, all groups of ELA-treated males consumed equal or less food than controls (**Fig. 1F**). In addition, excretion of energy via feces did not differ in any group relative to controls (**Extended Data Fig. 1D**), suggesting that increased adiposity cannot be explained by increased caloric intake or greater dietary energy harvest. To test whether increased adiposity was alternatively driven by lower energy expenditure, we used indirect calorimetry to measure energy expenditure (EE) in 30-week-old mice. We found reduced resting EE among L4 and HD males versus controls, with a similar trend in total EE (**Fig. 1I-J**), a result consistent with the lower lean mass and organ mass associated with ELA treatment.

In contrast, females proved resistant to the obesogenic effects of ELA treatment. Despite HD treatment inducing temporary growth stunting in both females and males, ELA-treated females of all dosages were similar to controls or trended in the opposite direction to that seen in males in every other measure of body size, body composition, and metabolic rate (**Extended Data Fig. 1E-M**). Whereas all groups of ELA-treated males had larger visceral fat deposits, L4 and HD females trended towards decreased visceral fat (**Extended Data Fig. 1I**). Whereas ELA-treated males exhibited reduced lean masses, organ sizes, and metabolic rates, females were indistinguishable from controls (**Extended Data Fig. 1H-M**).

Overall, both high-dose and low-dose ELA treatments promoted visceral adiposity in adult males at the expense of investment in lean tissue and organ mass that resulted in lower metabolic costs of tissue maintenance — characteristics that together imply a shift towards more frugal energy usage, or “metabolic thrift”^27^. Females were less phenotypically plastic in response to ELA treatment, exhibiting neither reductions in body maintenance costs nor subsequent increases in visceral adiposity as sequelae.

### High-dose and low-dose ELA elicit distinct changes in the gut microbiome

Recipients of all ELA treatments exhibited altered gut microbiota composition compared to age– and sex-matched controls, starting in gestation and for the duration of each treatment (**Fig. 2A-C, Extended Data Table 1**). Among ELA treatments, HD gut microbiomes differed compositionally from L4 and L30 microbiomes (PERMANOVA, P=0.001; **Fig. 2A**), were more compositionally distant from pre-treatment timepoints (mothers) and age– and sex-matched controls (offspring) during treatment (Wilcoxon rank-sum, P<0.01), and showed both delayed and incomplete post-treatment recovery of the gut microbiota compared to L4 microbiota (**Fig. 2B-C)**. HD treatment was characterized by up to 10,000-fold initial reductions in fecal bacterial density, whereas bacterial density under L4 and L30 treatments was minimally altered from controls (**Fig. 2D**). After controlling for variation attributable to treatment and litter, we did not detect any sex differences in the gut microbiota until 5 weeks of age (R^2^=0.008, p=0.009, PERMANOVA), after which sex differences persisted with increasing effect sizes that peaked at 20 weeks (R^2^=0.066, p=0.001) (**Extended Data Table 1**).

**Figure 2:**
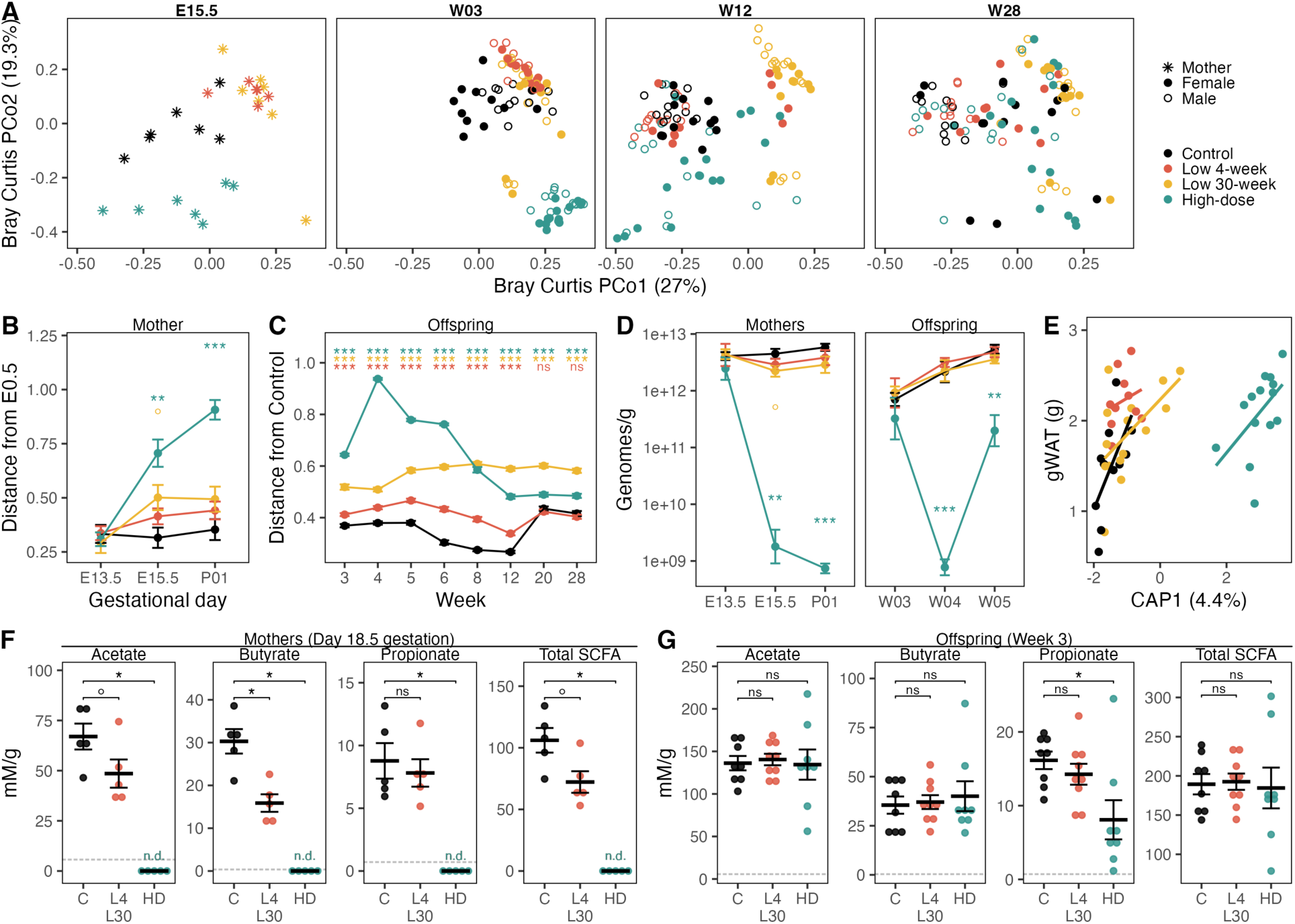
ELA alters gut microbiome composition and SCFA production in a dose-dependent manner. (**A**) Principal coordinate plots of fecal microbiome composition among pregnant mothers during treatment (E15.5) and among offspring at weeks 3, 12, and 28. (**B**) Bray-Curtis distances of the maternal fecal microbiome prior to treatment (E13.5), during treatment (E15.5), and 1 day post-partum (P01) versus the start of pregnancy (E0.5). (**C**) Bray-Curtis distances between offspring and age– and sex-matched control mice of different litters. (**D**) Fecal bacterial density in mothers and offspring as quantified by universal 16S qPCR. (**E**) Constrained analysis of principal coordinates (CAP) plot of gut microbial composition at week 4 and gWAT mass at week 28 among males, using the formula Bray-Curtis distance ∼ gWAT. (**F**) Normalized cecal short-chain fatty acid concentrations in pregnant dams at day 18.5 gestation and (**G**) in gestationally exposed offspring at 3 weeks of age. Dotted grey horizontal lines in F-G represent lower limit of detection for individual SCFAs. Data are mean ± s.e.m. Wilcoxon rank-sum test with control as reference group, ns = p≥0.1, ° = p<0.1, * = p<0.05, ** = p<0.01, *** = p<0.001.

Differential abundance analysis using MaAsLin2 identified several microbial taxa associated with ELA treatments (**Extended Data Table 2**), including reductions in *Allobaculum* and *Lactobacillus* (**Extended Data Fig. 2A-C**), which have been consistently reduced by ELA treatment in prior studies of ELA-induced obesity^9,11,14^. Constrained analysis of principal coordinates identified an axis of variation in early-life (week 4) but not adult (week 28) male gut microbiota that significantly correlated with the size of adult visceral fat deposits (PERMANOVA, P=0.054; **Fig. 2E**). This suggests that variable visceral adiposity among adult males might be related to interindividual variability in the gut microbial response to ampicillin treatment.

Perturbation of the gut microbiota led to disrupted production of SCFAs, which are known to play critical roles in energy homeostasis and metabolic programing during gestation^18,28^. By the end of gestation (E18.5), cecal concentrations of SCFAs in HD mothers were below detectable levels — a dramatic reduction compared with untreated mothers — whereas low-dose treatment (L4 or L30) elicited more modest reductions in acetate and butyrate but not propionate (**Fig. 2F**). Notably, the impacts of gestational HD treatment on cecal SCFA partially persisted between antibiotic pulses, with 3-week-old HD mice showing reduced cecal propionate (**Fig. 2G**). By 30 weeks of age, no reductions in cecal SCFAs were detectable in HD mice, although L4 males showed elevated cecal butyrate (**Extended Data Fig. 2D**). Combined with evidence of growth stunting during HD treatment, these data suggest that high-dose but not low-dose ampicillin treatment temporarily constricts energy availability during treatment at least partly through its impact on the gut microbiota.

### Plastic male phenotypic responses to ELA attenuate the negative effects of adult caloric restriction, an effect absent in less-plastic females

We next sought to test whether developmental plasticity prompted by energy-limiting ELA carries adaptive benefits that trade-off against consequences for metabolic health. We hypothesized that the observed developmental plasticity in males — including their smaller, more metabolically thrifty bodies — conferred fitness benefits under adult conditions of low energy availability. Likewise, we evaluated whether the apparent lack of metabolic thrift in ELA females came at the cost of fitness in other areas, such as reproduction or immunity, under conditions of low energy availability. To this end, we generated new litters of ELA-treated mice, focusing on the HD intervention because it induced stronger early-life energetic constraints and better represents the pulsed therapeutic antibiotic exposures of human infants. Starting at 8 weeks of age, HD and CON mice either continued with *ad libitum* feeding (AL) or were placed under 30% caloric restriction (CR), representing an adverse adult ecological exposure (**Fig. 3A**). This age was chosen because 8-week-old mice have completed the majority of their lean mass growth but do not yet exhibit any of the differences in visceral adiposity or organ mass observed at 30 weeks (**Fig. 1G, Extended Data Fig. 1N-O**). After 2 weeks of the dietary intervention, male and female mice were evaluated across a range of fitness parameters, including fertility and reproductive success, response to acute infection, body size, growth, and fat storage.

**Figure 3:**
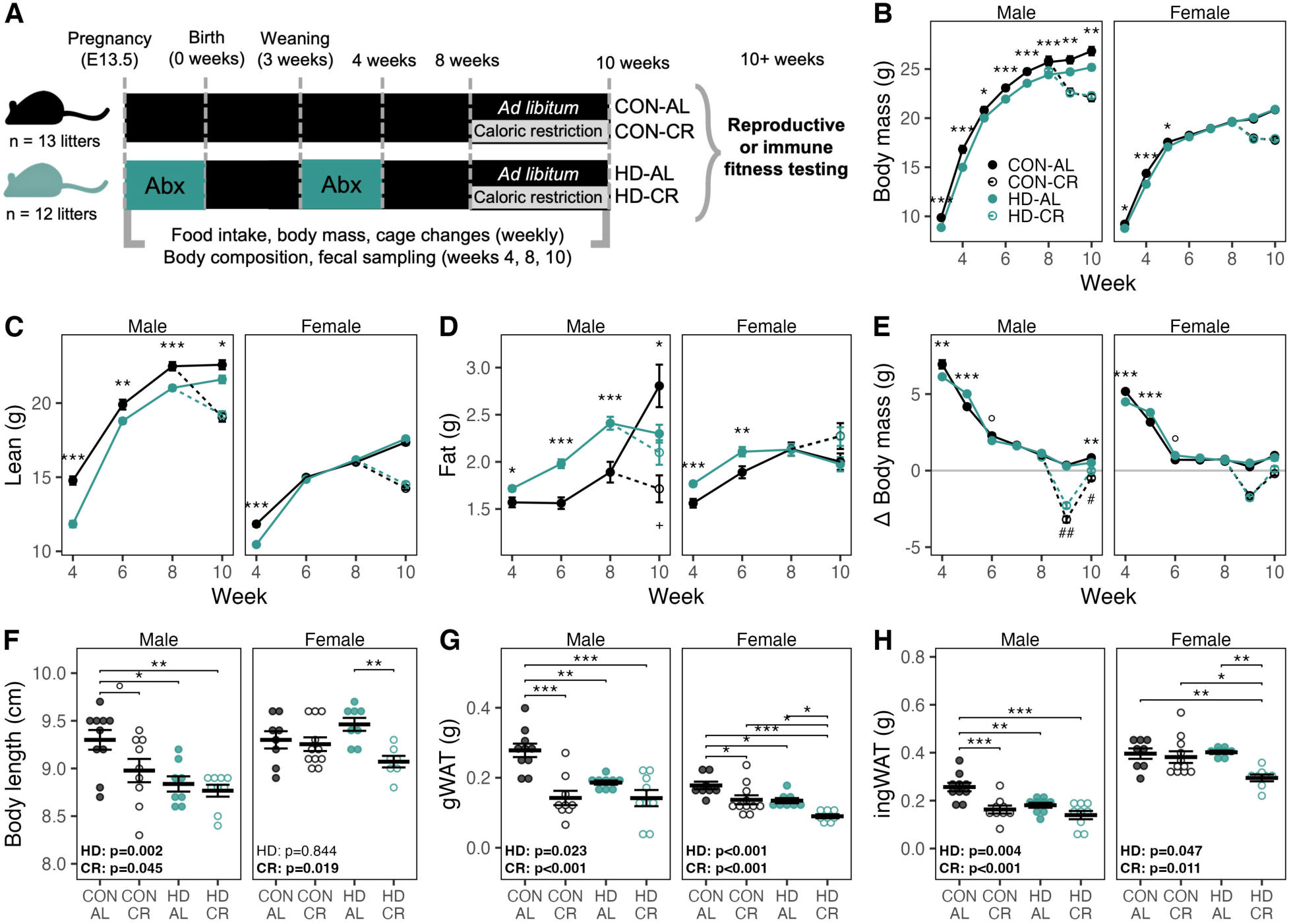
Growth and fat storage of ELA-treated mice under adult caloric restriction or *ad libitum* feeding. (**A**) Experimental design. (**B**) Body mass. (**C**) Lean body mass and (**D**) fat mass, as measured by EchoMRI. (**E**) Change in body mass from 3-10 weeks, with the y-axis value representing change observed at the week shown on the x-axis relative to the previous week. Endpoint (**F**) body length, (**G**) visceral fat as indexed by gonadal (epididymal or parametrial) white adipose tissue (gWAT) mass, and (**H**) subcutaneous fat as indexed by inguinal white adipose tissue (ingWAT), with the male endpoint at 10 weeks and the female endpoint at ∼13 weeks (immediately following pregnancy). Data are mean ± s.e.m. Statistical annotations for B-E: t-test: ° = p<0.1, * = p<0.05, ** = p<0.01, *** = p<0.001 for comparisons among AL mice, or ^+^ = p<0.1, ^#^ = p<0.05, ^##^ = p<0.01 for comparisons among CR mice. Statistical annotations for F-H: two-way ANOVA (∼ HD + CR), with significant features noted in bold, followed by Tukey’s HSD for pairwise comparisons: ° = p<0.1, * = p<0.05, ** = p<0.01, *** = p<0.001.

Replicating the growth phenotypes observed in our prior ELA experiments, HD treatment impacted body mass in young (≤5 weeks) male and female mice (**Fig. 3B**), reducing growth rates during the week of post-weaning HD treatment (**Fig. 3E**). Despite accelerated “catch-up” growth in HD mice of both sexes after cessation of antibiotics (**Fig. 3E**), HD males maintained their reduced body mass through 10 weeks of age (**Fig. 3B**), attributable to reduced lean mass (**Fig. 3C**). Unlike the pattern in males, catch-up growth in HD females was enough that HD females showed no alterations in growth, body size, or body composition at 10 weeks compared to CON females under AL feeding (**Fig. 3B-D**).

While young males and females both experienced the short-term energy-limiting effects of HD treatment, their divergent developmental responses led to starkly different outcomes under adult CR. Under CR, both HD and CON males lost body mass and particularly lean mass versus AL-fed controls (**Fig. 3B-E**). However, by 10 weeks of age, after 2 weeks of CR, CON-CR males had lost more weight and lean mass than HD-CR males (**Fig. 3C,E**). CR treatment also reduced body length at 10 weeks in CON-CR males relative to CON-AL males but had no effect on the body lengths of HD males (**Fig. 3F**). We found a similar pattern in visceral fat pad (gWAT) and subcutaneous fat pad (ingWAT) mass at 10 weeks, where the negative impact of CR observed among CON males was blunted in HD-CR males compared to vs HD-AL males (**Fig. 3G-H**). By contrast, whereas HD and CON females both lost body mass and lean mass under CR, there was no detectable difference between HD-CR and CON-CR groups in adult female growth rates, body mass, body, or body length (**Fig. 3B-F**).

These data suggest that metabolic plasticity by males in response to HD-ELA enables them to better tolerate adult conditions of energy limitation. Conversely, the lack of metabolic thrift by females in response to HD-ELA may help prevent them from developing obesity in an energy-rich adult environment but also provides no comparable buffer under conditions of adult energy limitation.

### ELA reduces female reproductive fitness in a manner exacerbated by caloric restriction

Life history theory predicts that differential costs of reproduction for male and females will favor sex-specific energetic trade-offs in response to energetic stress. We therefore hypothesized that while HD females showed few ELA-induced differences in energy allocation towards growth and maintenance, female reproduction might be especially sensitive to fluctuations in energy availability given the high energetic costs of mammalian gestation and lactation.

To directly quantify female reproductive fitness, we cohoused 10-week-old HD-AL, HD-CR, CON-AL or CON-CR females individually with an unexposed adult male for 10 days and tracked pregnancy outcomes and energy balance through birth (**Fig. 4A**). All pairings resulted in pregnancy and there were no detectable differences in the number of offspring, live or dead, associated with HD or CR treatment (**Fig. 4B**, **Extended Data Fig. 3A-B**). However, offspring birthweights showed additive decreases under both HD treatment and CR, with HD-CR mothers producing notably smaller offspring than all other groups (**Fig. 4C**). Therefore, while HD and CR treatments did not impact the total number of surviving offspring at birth, they did impact the predicted quality of each offspring, and the negative effects of CR were exacerbated for HD mothers.

**Figure 4:**
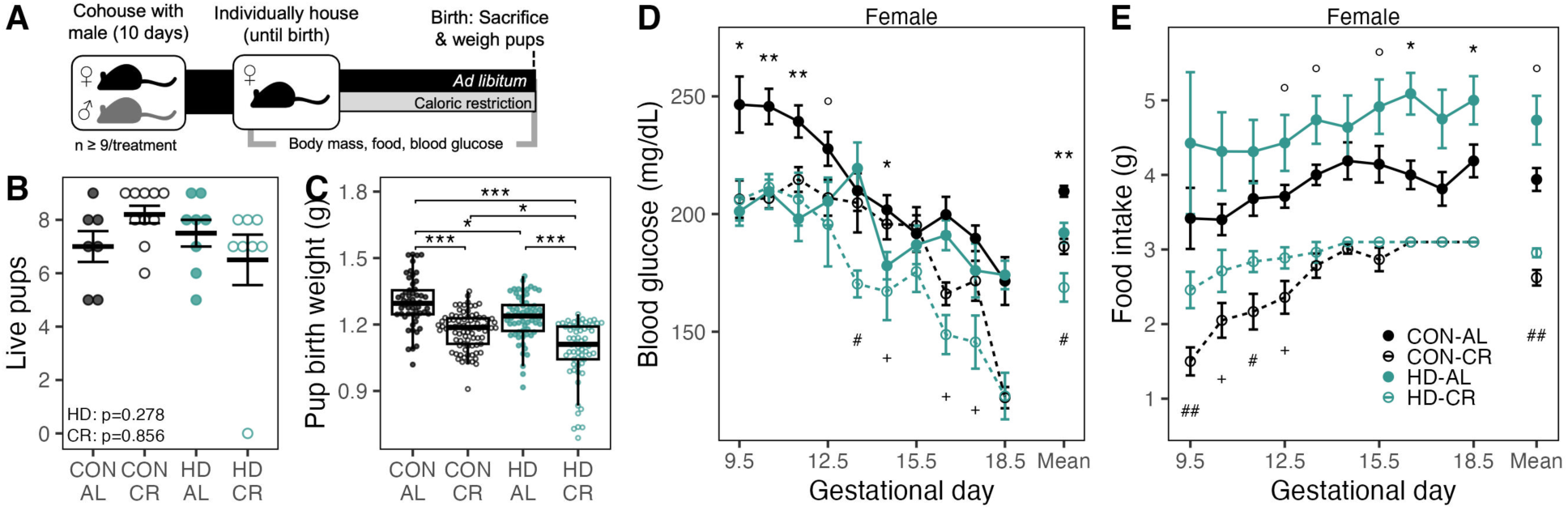
ELA and caloric restriction additively impair female reproductive fitness. (**A**) Experimental design of female reproductive fitness test. (**B**) Total live pups delivered per litter. (**C**) Birth weight of offspring. (**D**) Blood glucose and (**E**) food intake during pregnancy. Data are median ± IQR (B) or mean ± s.e.m (C-E). Statistical annotations for B: two-way ANOVA (∼ HD + CR); for C: linear mixed effects model of Pup mass ∼ ELA_status + Feeding + (1|Mother_ID), * = p<0.05, *** = p<0.001; for D-E: t-test: ° = p<0.1, * = p<0.05, ** = p<0.01, for comparisons among AL mice, or ^+^ = p<0.1, ^#^ = p<0.05, ^##^ = p<0.01 for comparisons among CR mice.

Smaller birth weights among the offspring of HD mothers were potentially related to greater pregnancy-related energetic stress. During the latter half of pregnancy, both HD-AL and HD-CR females ate more food and exhibited lower blood glucose levels than did CON females in the same feeding group (**Fig. 4D-E**). HD mothers ended pregnancy with smaller visceral gWAT fat stores than CON mice of the same feeding conditions, and subcutaneous ingWAT stores were also smaller among HD-CR mothers compared to CON-CR mothers (**Fig 3G-H**). Together, these data suggest that HD females experienced greater energy requirements during pregnancy and poorer energy transfer to their developing embryos.

To evaluate potential impacts of ELA on reproductive fitness in males, we measured testes size and performed epididymal sperm counts at 10 weeks. In contrast to females, we detected no differences in reproductive capacity associated with HD or CR treatment (**Extended data Fig. 3D-F**). Our results suggest that reproductive fitness in females is especially sensitive to energy limitation when combined with HD-ELA treatment, highlighting a lingering cost to female reproduction even two months after the cessation of antibiotic treatment. Thus, while males respond to the stress of ELA by reducing investment in lean mass, females may respond by reducing investment in reproductive capacity.

### ELA increases susceptibility to Campylobacter jejuni infection regardless of host sex or energy availability

Prior studies indicate that immune development is compromised by ELA-mediated effects on the gut microbiota^29^. We therefore tested whether the differential physiological responses to ELA treatment in males and females extended to differential susceptibility to infection under variable feeding conditions. We infected 10-week-old HD-AL, HD-CR, CON-AL and CON-CR mice of each sex with *Campylobacter jejuni* and monitored pathogen load and weight loss over the subsequent 3 days (**Fig. 5A**).

**Figure 5:**
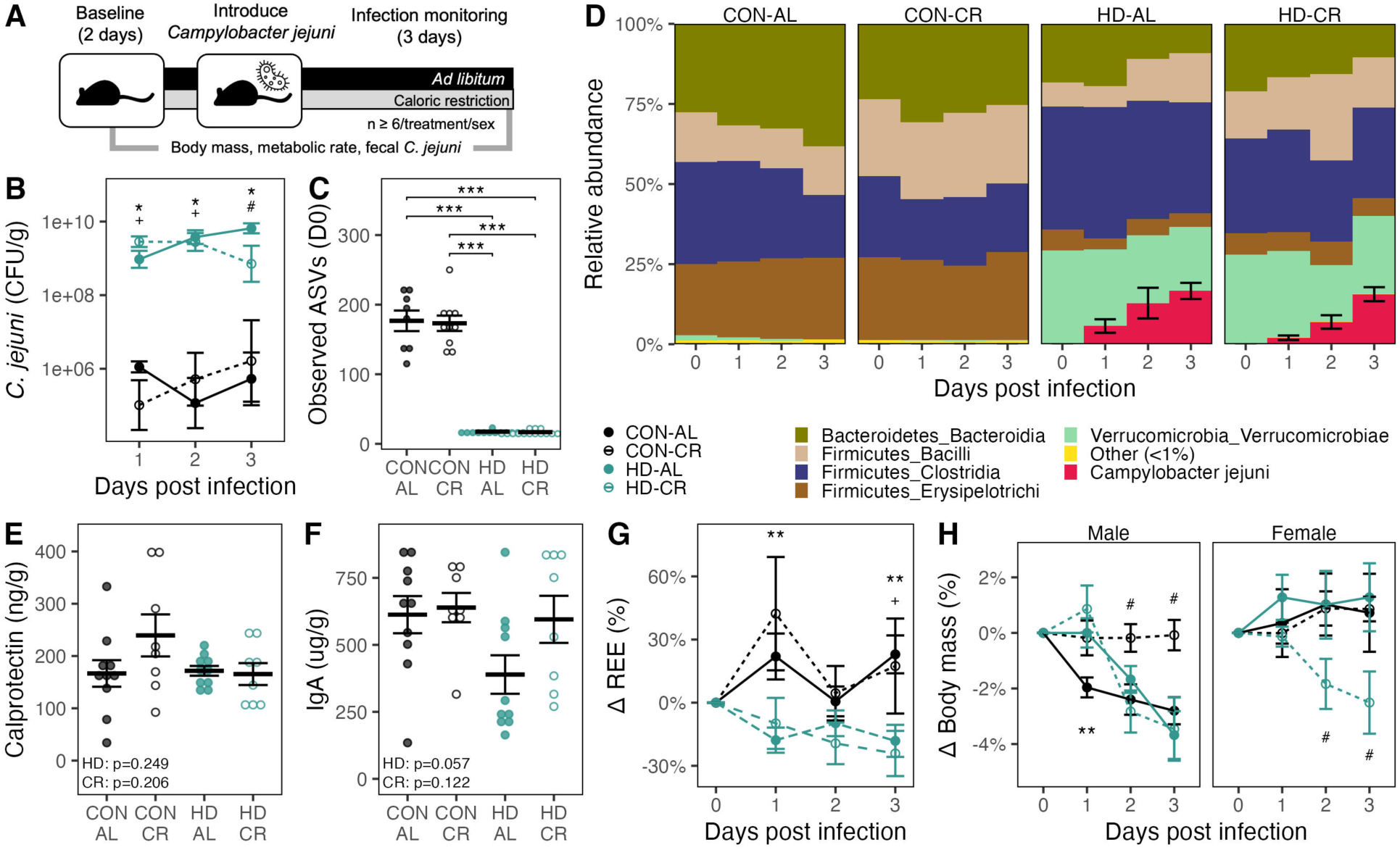
Outcomes of *Campylobacter jejuni* infection driven by ELA exposure. (**A**) Experimental design of immune fitness test. (**B**) Density of *C. jejuni* colony forming units (CFU) cultured from feces of infected mice. **(C**) Microbiome richness (count of observed ASVs) in feces immediately prior to *C. jejuni* infection. (**D**) Composition of fecal microbiome at baseline and during the 3-day immune challenge. (**E**) Fecal calprotectin, **(F**) fecal IgA, (**G**) change in resting energy expenditure (REE) from baseline, and (**H**) change in body mass from baseline during immune challenge. Data are mean ± s.e.m. Statistical annotations for B, G-H: Wilcoxon rank-sum test (B) or t-test (G-H) comparing among AL mice, * = p<0.05, ** = p<0.01 or among CR mice, ^+^ = p<0.1, ^#^ = p<0.05; for C: Wilcoxon rank-sum test, *** = p<0.001; for E-F: two-way ANOVA (∼ HD + CR).

Over the course of the infection period, HD mice of both sexes and feeding groups shed up to 10,000 times more live *C. jejuni* in feces than CON mice, with no additional effect of CR on live *C. jejuni* load in either HD or CON mice (**Fig. 5B**). Relative abundance of *C. jejuni* continued to increase over the duration of infection in HD mice (**Fig. 5D**), suggesting poor immunological control. High *C. jejuni* load in HD mice may have been partially driven by impaired colonization resistance, as HD gut microbiota contained <10% of the mean community richness found in CON mice prior to infection (**Fig. 5C**). Despite higher pathogen loads, HD mice did not exhibit greater inflammatory responses to *C. jejuni* infection than CON mice, regardless of feeding status, as evidenced by similar levels of fecal calprotectin and secretory IgA across treatments (**Fig. 5E-F**).

To test whether ineffective immune responses in HD mice were driven by energetic trade-offs, we examined RMR over the course of infection. Consistent with suppressed energy investment in the acute immune response of HD mice, both HD-AL and HD-CR mice exhibited decreases in resting energy expenditure (REE) over the course of infection relative to CON mice in the same feeding group (**Fig. 5G**). Infected HD mice also exhibited lower REE relative to their pre-infection baseline (HD-AL, p=0.079; HD-CR p = 0.021; Wilcoxon rank-sum test), an effect not observed in CON mice (CON-AL, p=0.740; CON-CR, p=1.00). While these fluctuations in REE could in theory arise from non-immune HD-mediated physiological changes (e.g., diet-induced or non-shivering thermogenesis), the strong fitness consequences of inefficient pathogen defense coupled with the absence of elevated markers of inflammation in HD mice suggest that ELA treatment constrains energy allocation to immune function. Further supporting the idea that ELA-induced energetic trade-offs constrain immune investment, HD treatment exacerbated the amount of weight lost by CR mice over the course of infection: by day 3, both male and female HD-CR mice had lost more body mass than CON-CR mice of the same sex (**Fig. 5H**).

Together, evidence of higher infection loads and suppressed acute immune response suggest that HD ELA treatment impaired immune development, with similarly negative consequences for male and female immune fitness.

### Male response to ELA is consistent with a predictive adaptive response to adverse adult environments, while female response promotes fitness in typical adult environments

To better understand the ultimate, evolutionary mechanisms driving the developmental response to ELA, we tested fitness outcomes of HD ELA against adaptive models for the developmental origins of obesity. Under the Predictive Adaptive Response hypothesis (PAR) — which posits that organisms with adverse developmental exposures can adapt to anticipated adverse adult environments — physiological responses that reduce fitness under resource-rich adult environments may allow individuals to succeed in adverse environments better than otherwise expected (**Fig. 6A**). Under the Developmental Constraints (DC) and Maternal Capital (MC) models — which posit that physiological responses to early-life adversity function to maximize survival during the adverse early-life period (DC) or as a maternal strategy to optimize lifetime reproductive success at the cost of the current offspring’s fitness (MC) — individuals that experienced adverse early-life conditions are expected to be disproportionately negatively affected by further adversity in adulthood (**Fig. 6A**). While all 3 models predict reduced fitness in ELA mice under AL feeding, PAR predicts the negative fitness impact of CR would be attenuated in ELA-CR versus CON-CR mice, whereas DC and MC predict the negative fitness impact of CR would be exacerbated in ELA-CR versus CON-CR mice.

**Figure 6:**
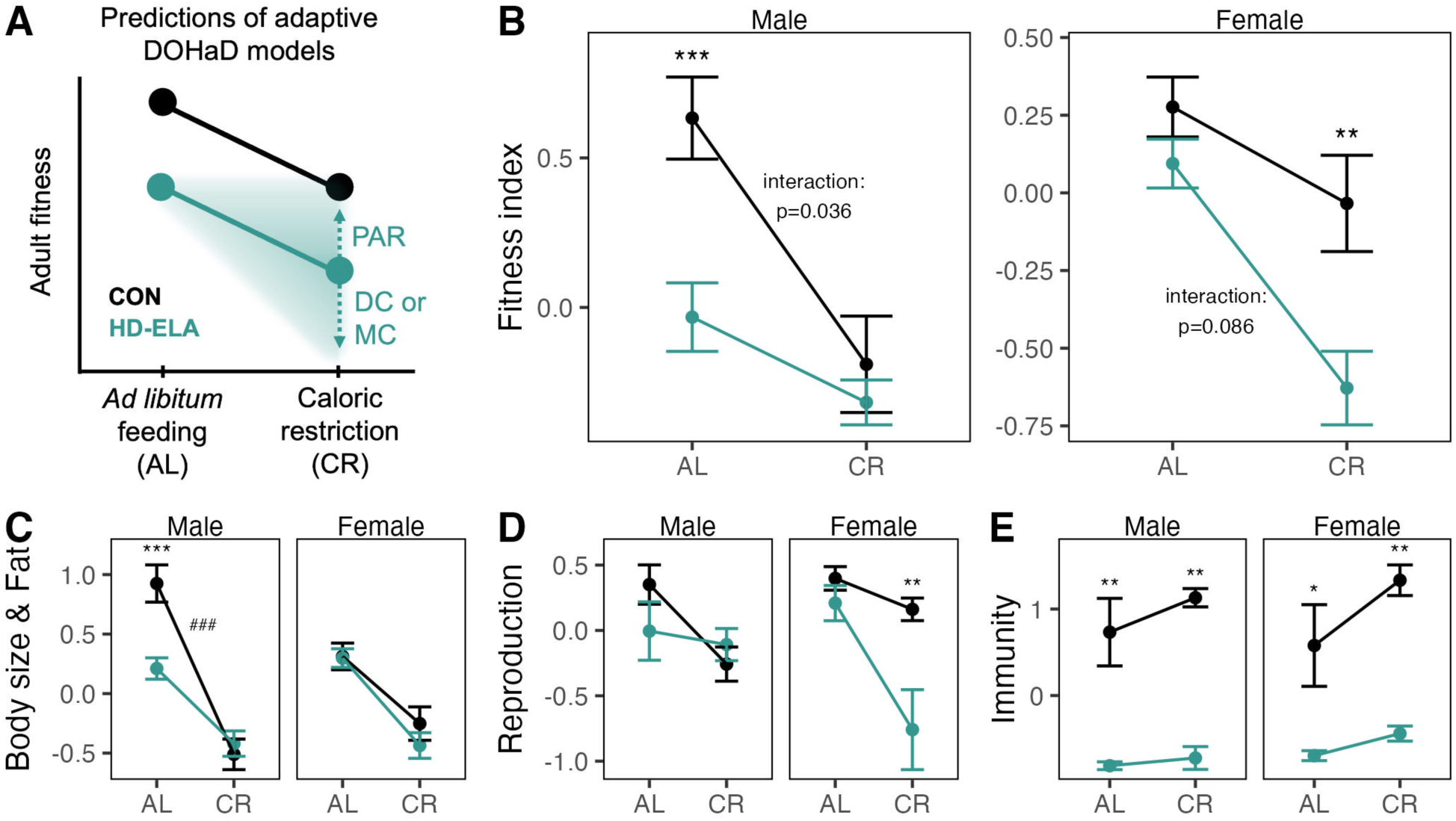
ELA elicits fitness trade-offs that differ by sex. (**A**) Predictions under adaptive developmental origins of health and disease models applied to the present study. (**B**) Composite fitness index of ELA mice under *ad libitum* feeding or caloric restriction. Components of fitness index related to (**C**) body size and fat storage, (**D**) reproduction, and (**E**) immunity. Data are mean ± s.e.m. Statistical annotations specify results of two-way ANOVA (∼ HD * CR) followed by pairwise t-tests, indicating significant differences between CON and HD, * = p<0.05, ** = p<0.01, *** = p<0.001; and significant interaction effects, ^###^ = p<0.001.

Given our observation that adult males and females experience the consequences of ELA treatment in different fitness currencies, we assessed these models using a combined index of fitness calculated as the per-mouse mean of Z-scores for body size and fat storage (body length, lean mass, total body fat mass, gWAT mass), reproduction/fertility [epididymal sperm counts and testes size (males) or live pup count and birth weights (females)], and acute immunity (*C. jejuni* load and weight loss at 3 days post-infection).

In males, our combined fitness index revealed a significant interaction effect of HD and CR treatment, with a reduction in fitness with HD compared to CON under AL feeding, but no difference under CR (**Fig. 6B**). In other words, HD males were better able than CON males to weather the adverse environment of CR, an outcome that is consistent with the PAR hypothesis but not with DC or MC. Considering the individual fitness components that comprise this index showed that the male pattern of PAR was driven largely by measures of body size and fat storage (**Fig. 6C**). Reproductive components of fitness were not affected by HD or CR in males (**Fig. 6D**), while immune fitness was reduced in all HD males but showed no significant interaction between HD and CR treatments (**Fig. 6E**). The male response to ELA therefore allows them to better maintain energy stores in an energy-restricted adult environment but is still generally disadvantageous for other aspects of fitness such as immunity.

In females, our combined fitness index indicated that ELA had limited effects on fitness under standard conditions of AL feeding but distinctly impaired fitness under CR (**Fig. 6B**), with the exacerbated impacts of CR most evident in the components related to reproduction (**Fig. 6D**). Under AL feeding, HD females showed impaired fitness relative to CON in measures related to immunity (**Fig. 6E**) and in specific fitness components such as gWAT (**Fig. 3G**) and pup birth weight (**Fig. 4C**). These differences were offset in the composite fitness index by the lack of detectable differences between HD-AL and CON-AL females in other metrics. Our data on ELA females therefore better fit the DC and MC models than PAR, but with the caveat that this does not apply to all aspects of fitness. We posit that females exhibit reduced developmental plasticity under HD ELA in order maximize fitness in a typical adult environment, but that this strategy incurs an additional fitness cost if they instead encounter an energy-limited adult environment.

## DISCUSSION

In this study, we showed that either pulsed high-dose or chronic low-dose exposure to ELA can promote obesity in males, but not females, via developmental changes characteristic of metabolic thrift, including reduced lean mass, organ size, and energy expenditure. As we previously hypothesized^3^, high-dose ampicillin treatment — but not low-dose treatment — impaired short-term energy harvest by the gut microbiota, characterized by stunted growth and reduced cecal SCFAs during treatment with more lasting reductions in cecal propionate. Low SCFA levels as a result of antibiotic treatment or germ-free status, particularly losses of propionate, have been shown to impair metabolic development and mediate adult risks of diet-induced obesity^18^. Additionally, the pattern we observed of growth stunting followed by more rapid catch-up growth after high-dose ampicillin treatment is one that also occurs under conditions of temporary food restriction in animal models and humans^22^, typically eliciting the same adaptations for metabolic thrift that we observed among male ELA-treated mice. It is therefore plausible that ELA-induced developmental changes in males may have been triggered by physiological mechanisms originally evolved to respond to early-life energy restriction.

Development of metabolic thrift in ELA-treated males likely helped them maintain body mass and fat stores when exposed as adults to environmental stress in the form of CR. The energetic buffering observed in ELA-treated males under adverse adult conditions is consistent with the PAR hypothesis, which predicts that developmental responses to adverse early-life exposures confer benefits in adverse adult environments. Although ELA-treated males experienced reduced impairments under CR, their fitness did not exceed that of controls under CR, suggesting that ELA treatment exacts a net fitness cost regardless of environmental conditions. Notably, PAR did not explain observed impacts on reproductive or immune fitness, as neither microbiome disruption in early life nor short-term caloric restriction induced detectable reductions in our indices of male fertility, and ELA treatment was markedly detrimental to immune fitness regardless of CR.

In striking contrast, females were resistant to the development of metabolic thrift and adult obesity under ELA, showing no significant long-term differences from controls in growth or body size under either low-dose or high-dose ELA or when combined with CR in adulthood. However, the reduced plasticity of females under ELA induced several trade-offs not evident in males. First, ELA-treated females had notably dysregulated metabolism during pregnancy, with signs of maternal hypoglycemia despite higher food consumption, as well as smaller pups. Additionally, females experienced exacerbated reductions in reproductive and immune fitness under CR. Jointly, these data suggest that maintenance of normal adult body size and composition by females under ELA comes at the cost of investment in reproduction and tolerance of energy-limited conditions.

Effects of ELA on female reproductive fitness highlight consequences of ELA that have previously been overshadowed by a focus on males and obesity outcomes. They also hint at wider, yet untested effects of ELA. For instance, our evidence of smaller offspring birthweights and maternal hypoglycemia prompts the hypothesis that ELA treatment may increase intergenerational risks of obesity or diabetes, as nutritional status during a mother’s own development can be as important as nutrition during pregnancy for offspring health^30^. Our results also raise the possibility that females exposed to ELA may experience more severe reproductive challenges than unexposed females, especially under conditions promoting negative energy balance such as food restrictive dieting. Such considerations are particularly pertinent to public health given widespread exposure to antibiotics in early life. Estimates based on US prescription rates indicate that children undergo an average of 3 courses of antibiotics before the age of 2^1^. Gestational and perinatal exposures to antibiotics are also extremely common, with routine administration of antibiotics to over 30% of pregnant women to prevent complications due to group B strep infection^31^ and the roughly 1/3 of mothers who deliver via cesarian section^32^.

The differential phenotypic and fitness responses that we observed in ELA-treated males and females illustrate both the importance of considering sex differences in metabolic disease research and the benefits of coupling proximate and ultimate lenses in investigating the etiology of metabolic disease. Sex differences in the developmental origins of metabolic diseases are well documented under a proximate lens, with mechanisms that include sex biases in gene imprinting and epigenetic modification, gene expression via X-chromosome inactivation, placental morphology and function, and the generally protective effect of higher estrogen levels^33,34^. Such proximate mechanisms help explain *how* dimorphic metabolic responses to early-life adversity arise, whereas ultimate mechanisms help explain *why* dimorphic metabolic responses to early-life adversity arise^3^. Through an ultimate lens, sex differences in the developmental origins of metabolic diseases might arise from the increased energetic burden of reproduction in females versus males, and therefore the comparatively greater payoffs to females of preserving metabolic capacity to support reproduction. Sexually dimorphic traits that are important for reproduction or competition for mates tend to exhibit increased sensitivity to environmental stressors^35^. For instance, among humans, lower socioeconomic status in childhood disproportionately affects height in males but pelvic width in females^35^. Our data follow a similar pattern, with ELA-induced reductions in body size and composition among males but reductions in the amount of energy directed to offspring during gestation in females.

Critically, our experiments were performed in laboratory mice and so we cannot yet infer whether similar trade-offs apply in humans exposed to ELA. Mice are in many ways an imperfect model for human health — for instance, they have less complex immune systems^36^, shorter life histories^37^, and more energetically intensive reproduction relative to body size^38^. Mice in laboratory settings may also experience delayed microbiome recovery after antibiotic treatment compared to free-living animals and humans that can benefit from microbial dispersal through social or environmental exposures^39^. Despite these challenges, our study does establish important similarities in the etiology of high-dose ELA-induced obesity in mice and humans. We observed a consistent male bias in the obesogenic and growth effects of ELA, a bias that has also been reported among humans^2^. Additionally, we found that increased visceral adiposity in ELA males was preceded by stunted growth, a characteristic that has also been observed in humans as a direct result of antibiotic perturbation of the gut microbiome^40^ and after bouts of diarrhea in young children^41^. Additional observational or epidemiological studies in humans will be required to elucidate the extent of parallels with our experimental studies in mice.

While further work in animal models and humans will continue to reveal the proximate and ultimate mechanisms underpinning ELA-induced obesity, we have shown that both pulsed high-dose and chronic low-dose ELA exposures promote visceral adiposity in males but not females via reduced investment in growth and metabolism. Additionally, we established contexts in which the physiological response of males to ELA and the attenuated response in females could be considered adaptive. To do this, we expanded upon previous studies that have explored the interaction between overnutrition (via high-fat diets) and ELA^9^ by examining ELA interactions with a model of undernutrition that is better positioned to illuminate fitness costs. In addition, we demonstrated that ELA impacts not only growth, metabolism, and immunity, but also previously hidden reproductive outcomes for females that affect offspring birth weights, raising the possibility of ELA-mediated intergenerational effects. Overall, our work establishes an adaptive framework for understanding interactions between health outcomes of ELA and environmental conditions that might be of particular interest to mitigating the impact of ELA in developing areas that not only experience increasing rates of obesity but also food insecurity and undernutrition.

## METHODS

### Animal housing and husbandry

To generate litters, 7-to-8-week-old male and female C57BL/6J mice were purchased from Jackson Laboratories and allowed to acclimate for ≥1 week. All females were mated with 1 of 10 breeder males that sired all litters in this study. Antibiotics were administered via drinking water (ampicillin, high-dose: 333 mg/L drinking water, ∼50 mg/kg body mass; low-dose: 6.7 mg/L, ∼1 mg/kg body mass) began at gestational day 13.5. Starting at weaning at 3 weeks of age, cages were changed weekly, and measurements were taken of offspring body mass and food intake (Food intake = Chow provided – Chow refused – Powdered chow in cage bottom). Fecal samples were collected from offspring starting at 3 weeks and every 1-4 weeks thereafter, with measurement of body composition via EchoMRI starting at 4 weeks and every 2-4 weeks thereafter. Unless otherwise specified, data reported concerns this F1 generation of mice, rather than their offspring (F2 generation) or the original breeding pair (F0 generation).

Mice were maintained on autoclaved PicoLab Mouse Diet 20 5058 provided *ad libitum* until the caloric restriction (CR) intervention or sacrifice. Mice subjected CR or their *ad libitum*-fed counterparts (AL) (**Fig. 3A**) were moved from group housing with same-sex littermates to individual housing at 8 weeks of age and half of the mice of each sex in each litter were put on 30% CR. Food was provided daily to mice in the CR group approximately 8 hours into the light period, with quantities scaled to body mass and based on previously observed food intake by age– and sex-matched controls [CR Food = 2.5 g + 0.1 × (Body mass – 25 g), with a minimum of 2.5 g for both males and females]. During this individual housing period from 8 to 10 weeks, food intake and body mass were measured either daily (CR) or semi-weekly (AL). After 10 weeks of age, mice were assigned for further profiling of either reproductive fitness or immune fitness.

Females undergoing reproductive fitness testing (**Fig. 4A**) were cohoused with a breeder male for 10 days, after which we measured body mass, food intake, and blood glucose daily until birth. Midway through the light period on the day of birth, dams and their offspring were weighed, sacrificed, and additional tissues and tissue weights were collected from the dam. For reproductive fitness testing in males, we performed epididymal sperm counts as a measure of fertility, following previously described methods^42^. For immune challenge testing, mice were transferred to autoclaved respirometry cages (see Metabolic rate measurement) and allowed 2 days to acclimate prior to introduction of *Campylobacter jejuni* (see *<u>Administration and detection of</u>* <u>Campylobacter jejuni</u>). Fecal samples and body mass measurements were collected daily over the 5-day period of acclimation and infection. Three days post-infection, mice were sacrificed, and various tissues were weighed and collected for further analysis.

All mice were housed in the specific pathogen-free Harvard University Biological Research Infrastructure facility with a 14:10 light/dark cycle, with all measurements and disbursement of food performed mid-way through the light period. Mouse work was performed under the supervision of the Harvard University Institutional Animal Care and Use Committee (Protocol #17-06-306).

### Metabolic rate measurement via indirect calorimetry

Mice were placed in a 3-cage open-flow indirect respirometry system consisting of Sable Systems Classic line equipment and Promethion line mouse cages (Sable Systems International, Las Vegas, NV). Mice were given ≥24 hours to acclimate to the respirometry cage prior to the start of measurement. Raw data were processed in R, where flow rates were corrected for standard temperature and pressure. Flow rates, % O_2_ and % H_2_O readings were then used to calculate oxygen consumption (VO_2_) and subsequently energy expenditure (EE) assuming an RQ of 0.8, as previously described^43^. To calculate resting EE over a 24-hour period, EE measurements were averaged over 5-min increments and the mean of the 3 lowest values was used.

### Cecal short-chain fatty acid quantification

Cecal samples were analyzed as described previously^14^. Briefly, 500 µl of HPLC-grade water was added to samples and vortexed for 10 min until homogenized. Samples were then centrifuged at 12,500 × g for 5 min, then 100 µl supernatant was collected for further processing. Supernatant pH of supernatant was adjusted to 2-3 by adding 10 µl of 50% sulfuric acid. Next, 10 µl of the internal standard (1% 2-methyl pentanoic acid solution) and 100 µl of anhydrous ethyl ether were added to each sample and vortexed for 2 min before centrifugation at 5,000 × g for 2 min.

The upper ether layer was transferred into an Agilent sampling vial for analysis using an Agilent 7890B system with flame ionization detector (FID) (Agilent Technologies, Santa Clara, CA) as previously described^44^. Briefly, a high-resolution gas chromatography capillary column 30 m x 0.25 mm coated with 0.25 µm film thickness (DB-FFAP) was used for the detection of volatile acids (Agilent Technologies, Santa Clara, CA) with oven temperature at 145°C and the FID and injection port set to 225°C and nitrogen as the carrier gas. 5 µl of extracted sample was injected for analysis and the run time was 11 min. Chromatograms and data integration were carried out using the OpenLab ChemStation software (Version C01.07Agilent Technologies, Santa Clara, CA). For the standard solution, a volatile acid mix containing 10 mM of acetic, propionic, isobutyric, butyric, isovaleric, and valeric acids was used (CRM46975, Sigma-Aldrich St. Louis, MO). A standard stock solution containing 1% 2-methyl pentanoic acid (Sigma-Aldrich St. Louis, MO) was prepared as an internal standard control for the volatile acid extractions. Lower limits of detection for each acid were as follows: 1.25 mM (acetate), 0.156 mM (propionate), 0.078 mM (butyrate, isobutryate, valerate, isovalerate).

### Fecal energy excretion

Used cage bottoms from Week 24 were collected and sifted to collect total fecal production. Approximately 1.8 g of the collected feces were then combusted in a bomb calorimeter (Parr Instrument Co., 6050 Calorimeter) to determine the energy density of each sample. Total energy excretion was calculated as Total fecal production × Fecal energy density.

### Oral glucose tolerance tests

At 26 weeks of age, mice were fasted for approximately 6 hours, then given a dose of 1.5 mg glucose/g body mass, delivered via oral gavage. Blood glucose was measured from tail blood using a Clarity BG1000 blood glucometer (Clarity Diagnostics) at 0, 15, 30, 60, 90, and 120 min following glucose administration. All blood glucose measurements were taken in duplicate with additional replicate measurements taken in the case of high variability between replicates (≥20 mg/dL).

### 16S rRNA gene sequencing

DNA was extracted from mouse fecal samples using E.Z.N.A. Soil DNA Kit (Omega BioTek, D5625-00S) following manufacturer’s instructions. Next, 16S rRNA gene was PCR-amplified using custom-barcoded 515F and 806R primers targeting the V4 region of the gene. PCR was performed on each sample in triplicate with sample-specific negative controls with the following protocol: 95°C for 3 min; 35 cycles of 94°C for 45 s, 50°C for 30 s, and 72°C for 90 s; and a final extension at 72°C for 10 min. We then quantified samples with Quant-iT Picogreen dsDNA Assay Kit (Invitrogen) prior to pooling samples evenly by DNA content. The resulting 16S rDNA pools were cleaned with AmpureXP beads (Agencourt). The cleaned pools were then loaded onto a mini 1.5% agarose gel, run for 50 min at 50V, and sequences of 381 bp length were re-extracted using the Qiaquick Gel Extraction Kit (Qiagen). All pools underwent 2×150 bp sequencing across 3 lanes of an Illumina NovaSeq platform.

Sequences were processed in QIIME2^45^, first by de-noising with Dada2 and truncating at 149 bp to ensure maximum sequence quality, resulting in read depths of 342,517 ± 112,208 across the n=1,037 samples associated with the 30-week study and read depths of 99,351 ± 69,183 across the n=707 samples associated with the 10-week study and immune challenge. Only forward reads were used for analysis. Taxonomy was assigned using the GreenGenes classifier^46^. The taxonomy and ASV feature tables were then imported into R (version 4.3.2) using qiime2R (version 0.99.6) and processed using phyloseq (version 1.46.0)^47^. First, each sample was pruned of very low abundance ASVs, defined as ≤3 reads per study pool. For the 30-week study, reads were subsampled evenly at 90,000 reads/sample which excluded 4 samples with <20,000 reads, resulting in n=1,033 samples. For the 10-week study, reads were subsampled at 20,000 reads/sample which excluded 4 samples, resulting in n=703 samples. Further processing of 16S rRNA gene sequences was then performed in R using phyloseq for calculation of distance matrices and ordinations, vegan (version 2.6-4) for PERMANOVA tests, and MaAsLin2 (version 1.16.0) for identifying differentially abundant taxa using general linear models.

### qPCR of the 16S rRNA gene

Quantitative PCR of the universal bacterial 16S rRNA gene was performed as described previously^14^. Briefly, we performed qPCR on the V4 region of the 16S rRNA gene (515F and 806R) in triplicate, using a standard curve on each plate based on genomic DNA isolated from a pure culture of *Escherichia coli* (ATCC 47076). We used the following recipe for each PCR reaction: 12.5 μl SYBR Green qPCR mix, 2.25 μl of each non-barcoded primer (515F and 806R), 6 μl nuclease-free water, and 2 μl template DNA, for a total volume of 25 μl per well. We ran the qPCR reactions on a BioRad CFX 96-well Real-Time PCR thermocycler with the following protocol: initial denature at 94°C for 15 min; 40 cycles of 95°C for 15 s, 50°C for 40 s, and 72°C for 30 s. To calculate 16S rRNA gene abundance, we first multiplied DNA concentrations of each sample as measured via qPCR then divided by the mass of the original fecal sample and multiplied by 2.03 x 10^5^ — an estimate of genome-equivalents per ng DNA based on a mean gut microbial community genome size of 4.50 Mbp^48^.

### Administration and detection of Campylobacter jejuni

*C. jejuni* (ATCC BAA-2151) was grown in brain heart infusion broth (BHI) (BD 214010) and incubated for 18-24 hours at 37°C in the GasPak EZ CampyPouch System (BD 260685). *C. jejuni* cultures were concentrated via centrifugation at 1,000 × g for 5 min and normalized to an optical density at 600 nm (OD_600_) of 0.2 (∼2 × 10^8^ CFU/ml) before administration. Mice were inoculated with 150 µl of the concentrated normalized suspension via oral gavage. To confirm OD_600_ estimates of *C. jejuni* density in the inoculum and quantify live *C. jejuni* shedding in the feces of infected mice, we performed 10-fold serial dilutions (×7) of the inoculum or a fecal homogenate (1:50 feces to PBS) and plated these on *C. jejuni*-selective plates (Hardy Diagnostics G339) that were incubated for ∼48 hours in the GasPak EZ CampyPouch System at 37°C until colonies were easily distinguishable. Culture based quantification was confirmed qPCR as described above (see *<u>qPCR of the 16S rRNA gene</u>*), but with *Campylobacter* specific primers pairs (F: CTGCTAAACCATAGAAATAAAATTTCTCAC, R: CTTTGAAGGTAATTTAGATATGGATAATCG)^49^ and a standard curve composed of genomic DNA isolated from a pure culture of pure *C. jejuni*.

### Quantification of fecal calprotectin and secretory IgA

We quantified calprotectin and secretory IgA (sIgA) in fecal samples from mice 3 days after *C. jejuni* infection by ELISA using commercially available kits: Mouse / Rat Calprotectin ELISA (Alpco 30-6936) and Mouse IgA ELISA Kit (Bethyl Laboratories E99-103). Prior to quantification, fecal samples were diluted 100-fold for calprotectin quantification and 1,000-fold for sIgA quantification, so that expected levels fell within the range of detection for each kit. Both assays were then performed according to manufacturer instructions.

### Fitness index calculations

To consider overarching effects of ELA across different measures of fitness, we generated a composite index capturing metabolic, reproductive, and immune outcomes. The metabolic component of the fitness index considered lean body mass, body length and total fat mass at 10 weeks, and mass of gWAT deposits at sacrifice. The reproductive component of the fitness index for females considered average pup mass and the number of live pups whereas the reproductive component of the fitness index for males captured epididymal sperm density and testes mass at 10 weeks. The immune component of the fitness index considered *Campylobacter jejuni* load (-logCFU, reversed so that higher values represent higher fitness) and the change in body mass between baseline and day 3 post-infection.

Because all mice had associated data on body size and composition at 10 weeks but could only undergo one of the subsequent fitness tests (reproductive or immune testing, not both), fitness scores for each mouse reflected 2X weighting of reproductive and immune metrics to avoid unintentional overrepresentation of body size and composition in the fitness index. The composite fitness index was therefore calculated as follows: *Fitness index* = [Z_score(*each metabolic component*) + 2 × Z_score(*each immune or reproductive component*)] / Ι.*(number of components considered)*.

### Statistical analysis

For simple comparisons between group means, we used Kruskal-Wallis tests followed by Wilcoxon rank-sum tests using the control group as a reference. To identify the combined effects of ELA (HD vs CON) and feeding status (CR vs AL), we used two-way ANOVA tests to compare differences in group means by each treatment type, followed by Tukey’s Honest Significant Difference test for pairwise comparisons. For specific comparisons within treatment variables (HD vs CON within CR or AL groups, and vice versa), t-tests or Wilcoxon rank-sum tests were uses as appropriate based on normality of data distribution. The Holm-Sidak method was used to correct for multiple comparisons. Some data types such as food intake and pup birth weight were analyzed using linear mixed effects (LME) models in order to control for the random effects of timepoint (food intake) and litter ID (pup birth weight). Unless otherwise noted, figure annotations show only significant (p≤0.05) or marginally significant (p≤0.01) comparisons.

We used PERMANOVA (permutational analysis of variance) tests with 999 permutations to assess variation in Bray-Curtis distances among gut microbiota across multiple variables using vegan (version 2.6-4). To identify genera that varied significantly with ELA treatment, we used LME models analyzed by MaAsLin2 with the following terms: fixed effects: Treatment; random effects: MouseID, LitterID, Timepoint. All MaAsLin2 models employed false discovery rate (FDR) corrections for multiple comparisons. All statistical analyses were conducted in R (version 4.2.3) using tidyverse packages (version 2.0.0).

## ACKNOWLEDGMENTS

We thank Martin Blaser, Daniel Lieberman, Terence Capellini, and members of the Carmody Lab for feedback on the project and manuscript. We also thank the staff of Harvard Office of Animal Resources, Harvard Bauer Core Facility, and the Massachusetts Host-Microbiome Center for animal care, sequencing, and SCFA quantification services and assistance, respectively. This study was supported by a National Science Foundation Graduate Research Fellowship (to L.D.S.) and awards from The William F. Milton Fund (to R.N.C.), Harvard Dean’s Competitive Fund for Promising Scholarship (to R.N.C.), and National Science Foundation (BCS-2142073 to L.D.S. and R.N.C.).

## AUTHOR CONTRIBUTIONS

L.D.S. and R.N.C conceived of the project and designed the study. L.D.S. performed the initial 30-week mouse experiment. L.D.S, G.R., and E.C. performed the subsequent mouse experiments and prepared gut effluent samples for 16S rRNA gene sequencing. L.D.S. and E.C. cultured, administered, and quantified *C. jejuni*. L.D.S. performed all other assays (metabolic rate measurements, fecal energy excretion quantification, qPCR, ELISAs) and conducted all data analysis. L.D.S. and R.N.C. interpreted the data and drafted the manuscript, with feedback from all authors. R.N.C. supervised the project.

## DATA AVAILABILITY

The 16S rRNA gene sequencing data described in this study is deposited to the NCBI Sequence Read Archive under submission number SUB15554908. The R code used in data analysis will be made available upon request.

