## Supplemental materials for "Early-life microbiota disruption by antibiotics elicits fitness trade-offs that differ by sex"

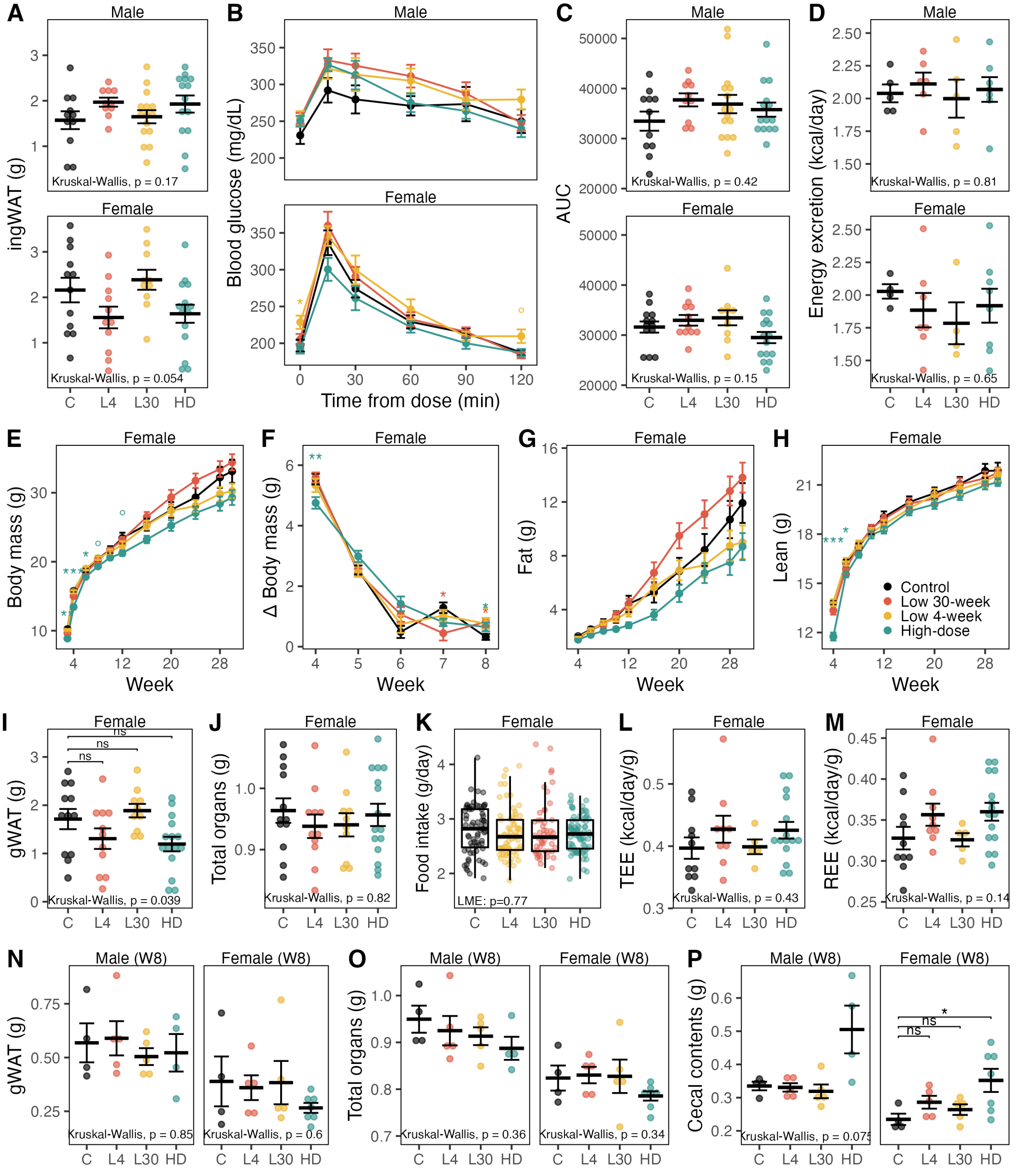


**Extended Data Figure 1: Phenotypic outcomes of low- and high-dose ELA treatment in males and females.**

(**A**) Mass of inguinal fat pad (ingWAT) at 30 weeks. (**B**) Blood glucose during oral glucose tolerance test at 26 weeks of age and (**C**) area under the curve of blood glucose timecourse. (**D**) Mean daily fecal energy excretion at 24 weeks. (**E**) Body mass from 3 to 30 weeks. (**F**) Change in body mass from 3-8 weeks, with the y-axis value representing change observed at the week shown on the x-axis relative to the previous week. (**G**) Fat mass and (**H**) lean body mass from 4 to 30 weeks, as measured by EchoMRI. (**I**) Mass of gonadal (perimetrial) white adipose tissue (gWAT), a visceral fat deposit, at

30 weeks of age. (**J**) Sum of organ masses (heart, spleen, kidney, gonads, brain) at 30 weeks. (**K**) Mean daily food intake per mouse. (**L**) Total energy expenditure (TEE) and (**M**) resting energy expenditure (REE), as measured by indirect calorimetry. (**N**) gWAT mass, (**O**) sum or organ masses (heart, spleen, kidney, brain), and (**P**) mass of cecal contents of mice sacrificed at 8 weeks of age. Data are mean ± s.e.m. Statistical annotations for A-J and L-P: Krukall-Wallis tests with significant results followed by Wilcoxon rank-sum test with control as reference group. For K, linear mixed effects model (LME) of Food intake ~ Treatment + (1|Cage_ID) + (1|Timepoint). Significance levels: ns = $p \geq 0.1$, ° = $p<0.1$, * = $p<0.05$, ** = $p<0.01$, *** = $p<0.001$.

| Week | Terms | Treatment Overall | | High-dose v Control | | Low 4-week v Control | | Low 30-week v Control | | Sex | | Treatment*Sex | | Litter | |
|---|---|---|---|---|---|---|---|---|---|---|---|---|---|---|---|
| | | $R^2$ | *P* | $R^2$ | *P* | $R^2$ | *P* | $R^2$ | *P* | $R^2$ | *P* | $R^2$ | *P* | $R^2$ | *P* |
| 3 | Treatment + Litter + Sex | 0.344 | **0.001** | 0.341 | **0.001** | 0.125 | **0.001** | 0.116 | **0.001** | 0.005 | 0.137 | | | 0.458 | **0.001** |
| 3 | Treatment * Sex | 0.344 | **0.001** | 0.341 | **0.001** | 0.125 | **0.001** | 0.116 | **0.001** | 0.011 | 0.141 | 0.019 | 0.582 | | |
| 4 | Treatment + Litter + Sex | 0.485 | **0.001** | 0.456 | **0.001** | 0.217 | **0.001** | 0.158 | **0.001** | 0.003 | 0.553 | | | 0.264 | **0.001** |
| 4 | Treatment * Sex | 0.485 | **0.001** | 0.456 | **0.001** | 0.217 | **0.001** | 0.158 | **0.001** | 0.003 | 0.658 | 0.010 | 0.904 | | |
| 5 | Treatment + Litter + Sex | 0.457 | **0.001** | 0.488 | **0.001** | 0.212 | **0.001** | 0.261 | **0.001** | 0.008 | **0.009** | | | 0.368 | **0.001** |
| 5 | Treatment * Sex | 0.457 | **0.001** | 0.488 | **0.001** | 0.212 | **0.001** | 0.261 | **0.001** | 0.010 | 0.093 | 0.022 | 0.161 | | |
| 6 | Treatment + Litter + Sex | 0.501 | **0.001** | 0.586 | **0.001** | 0.152 | **0.001** | 0.355 | **0.001** | 0.012 | **0.003** | | | 0.323 | **0.001** |
| 6 | Treatment * Sex | 0.501 | **0.001** | 0.586 | **0.001** | 0.152 | **0.001** | 0.355 | **0.001** | 0.010 | 0.077 | 0.014 | 0.572 | | |
| 8 | Treatment + Litter + Sex | 0.336 | **0.001** | 0.243 | **0.001** | 0.140 | **0.001** | 0.421 | **0.001** | 0.008 | **0.031** | | | 0.463 | **0.001** |
| 8 | Treatment * Sex | 0.336 | **0.001** | 0.243 | **0.001** | 0.140 | **0.001** | 0.421 | **0.001** | 0.018 | **0.033** | 0.011 | 0.938 | | |
| 12 | Treatment + Litter + Sex | 0.327 | **0.001** | 0.160 | **0.001** | 0.020 | 0.082 | 0.353 | **0.001** | 0.028 | **0.001** | | | 0.459 | **0.001** |
| 12 | Treatment * Sex | 0.327 | **0.001** | 0.160 | **0.001** | 0.020 | 0.410 | 0.353 | **0.001** | 0.031 | **0.005** | 0.023 | 0.319 | | |
| 20 | Treatment + Litter + Sex | 0.188 | **0.001** | 0.052 | **0.001** | 0.013 | 0.326 | 0.204 | **0.001** | 0.066 | **0.001** | | | 0.489 | **0.001** |
| 20 | Treatment * Sex | 0.188 | **0.001** | 0.052 | **0.008** | 0.013 | 0.674 | 0.204 | **0.001** | 0.090 | **0.001** | 0.041 | **0.040** | | |
| 28 | Treatment + Litter + Sex | 0.167 | **0.001** | 0.055 | **0.001** | 0.025 | **0.046** | 0.193 | **0.001** | 0.056 | **0.001** | | | 0.466 | **0.001** |
| 28 | Treatment * Sex | 0.167 | **0.001** | 0.055 | **0.008** | 0.025 | 0.292 | 0.193 | **0.001** | 0.077 | **0.001** | 0.030 | 0.210 | | |

**Extended Data Table 1: PERMANOVA tests of gut microbiome composition under low-dose or high-dose ELA treatment.**
Results from two different models, including either a Treatment by Sex interaction term or additional Litter effect. Data are subset first by each week, then by each treatment pair. PERMANOVA tests were run for each data subset, with the effects listed for Sex, Treatment*Sex, and Litter taken from the test run on all treatments together.

See separate excel file
**Extended Data Table 2: Microbial biomarkers of ELA treatment identified using MaAslin2.**

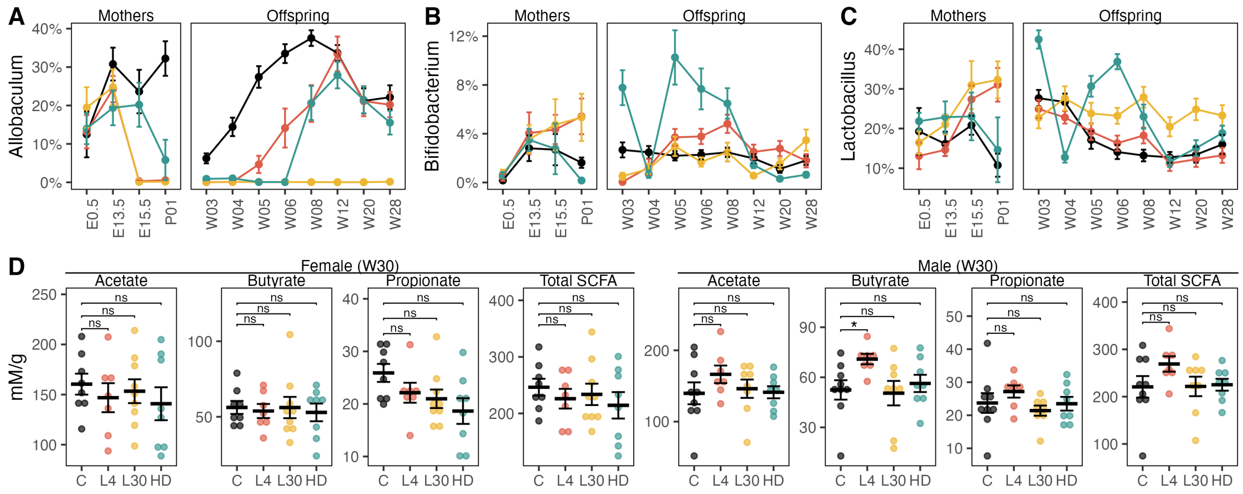


**Extended Data Figure 2: Gut microbiome changes under ELA treatment.**
(**A-C**) Relative abundances of fecal microbial genera that were identified by Maaslin2 as differing significantly from control under one or more ELA treatment regimens. (**D**) Cecal short-chain fatty acid (SCFA) concentrations in ELA treated mice at 30 weeks of age. Wilcoxon rank-sum test with control as reference group, ns = p≥0.1, * = p<0.05.

| | CAP1 | Phylum | Order | Family | Genus | Species | C | HD | L4 | L30 |
|---|---|---|---|---|---|---|---|---|---|---|
| Negative | -0.853 | Firmicutes | Lactobacillales | Lactobacillaceae | Lactobacillus | Unclassified | 27.447% | 0.908% | 23.954% | 29.266% |
| | -0.230 | Firmicutes | Erysipelotrichales | Erysipelotrichaceae | Allobaculum | Unclassified | 10.628% | 0.675% | 0.104% | 0.074% |
| | -0.186 | Bacteroidetes | Bacteroidales | S24-7 | Unclassified | Unclassified | 9.056% | 0.320% | 10.707% | 8.156% |
| | -0.160 | Bacteroidetes | Bacteroidales | S24-7 | Unclassified | Unclassified | 8.672% | 0.309% | 10.230% | 9.841% |
| | -0.154 | Bacteroidetes | Bacteroidales | S24-7 | Unclassified | Unclassified | 12.391% | 0.286% | 14.325% | 8.662% |
| | -0.098 | Bacteroidetes | Bacteroidales | S24-7 | Unclassified | Unclassified | 4.688% | 0.142% | 5.269% | 5.773% |
| | -0.082 | Firmicutes | Turicibacterales | Turicibacteraceae | Turicibacter | Unclassified | 6.345% | 0.074% | 0.028% | 0.021% |
| | -0.065 | Firmicutes | Clostridiales | Clostridiaceae | Unclassified | Unclassified | 4.963% | 0.026% | 0.010% | 0.009% |
| | -0.047 | Actinobacteria | Bifidobacteriales | Bifidobacteriaceae | Bifidobacterium | pseudolongum | 2.109% | 0.106% | 2.291% | 0.838% |
| | -0.033 | Bacteroidetes | Bacteroidales | S24-7 | Unclassified | Unclassified | 1.780% | 0.070% | 2.350% | 2.617% |
| Positive | 0.382 | Proteobacteria | Pseudomonadales | Pseudomonadaceae | Pseudomonas | Unclassified | 0.000% | 18.859% | 0.000% | 0.000% |
| | 0.203 | Firmicutes | Clostridiales | Unclassified | Unclassified | Unclassified | 0.011% | 5.322% | 0.024% | 0.031% |
| | 0.129 | Firmicutes | Lactobacillales | Lactobacillaceae | Lactobacillus | helveticus | 0.000% | 4.256% | 0.000% | 0.001% |
| | 0.088 | Bacteroidetes | Bacteroidales | S24-7 | Unclassified | Unclassified | 0.003% | 0.012% | 0.002% | 2.238% |
| | 0.080 | Firmicutes | Lactobacillales | Streptococcaceae | Streptococcus | Unclassified | 0.002% | 8.761% | 0.002% | 0.002% |
| | 0.069 | Bacteroidetes | Bacteroidales | S24-7 | Unclassified | Unclassified | 0.001% | 0.008% | 0.001% | 1.610% |
| | 0.042 | Firmicutes | Lactobacillales | Lactobacillaceae | Lactobacillus | Unclassified | 0.000% | 1.612% | 0.000% | 0.000% |
| | 0.039 | Firmicutes | Bacillales | Staphylococcaceae | Staphylococcus | succinus | 0.014% | 7.058% | 0.003% | 0.001% |
| | 0.037 | Firmicutes | Lactobacillales | Streptococcaceae | Lactococcus | Unclassified | 0.000% | 1.326% | 0.000% | 0.000% |
| | 0.037 | Bacteroidetes | Bacteroidales | S24-7 | Unclassified | Unclassified | 0.001% | 0.000% | 0.000% | 0.887% |

**Extended Data Table 3: Components of constrained analysis of principal coordinate analysis.** Showing the top 20 ASV components of the CAP1 axis, which correlates gut microbiota composition at 4 weeks to visceral adiposity (gWAT mass) at 30 weeks in males. The right four columns indicate mean relative abundance of each ASV in each treatment group.

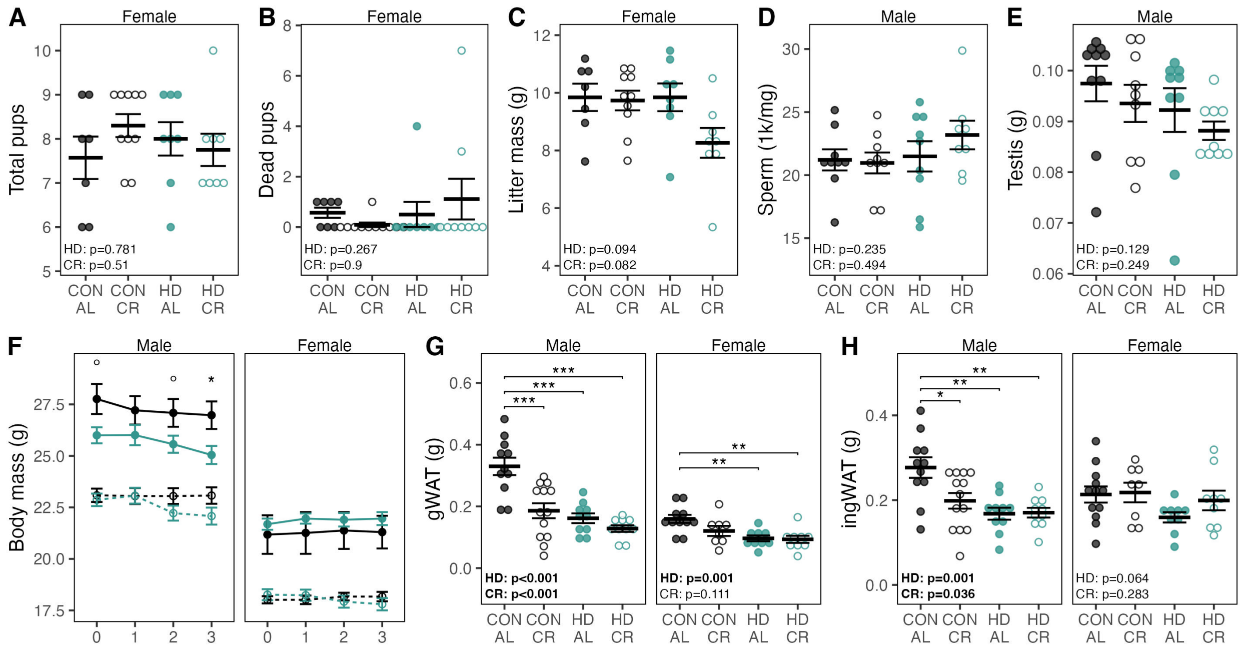


**Extended Data Figure 3: Additional outcomes from reproduction and immune fitness testing.** (**A**) Total pups delivered per litter (live and dead). (**B**) Total dead pups delivered per litter. (**C**) Total mass of litter. (**D**) Epididymal sperm counts and (**E**) mass of right testis at 10 weeks. (**F**) Body mass trajectory during *Campylobacter jejuni* challenge. (**G**) Gonadal/perimetrial fat pad (gWAT) mass and (**H**) inguinal fat pad (ingWAT) mass at conclusion of *Campylobacter jejuni* challenge (day 3). Data are mean ± s.e.m. Statistical annotations for A-D, G-H: two-way ANOVA (~ HD + CR), with significant features noted in bold, followed by Tukey's HSD for pairwise comparisons: * = $p<0.05$, ** = $p<0.01$, *** = $p<0.001$. Statistical annotations for F: t-test: ° = $p<0.1$, * = $p<0.05$ for comparisons among AL mice, with no significant differences among CR mice.
